# A maize male gametophyte-specific gene encodes ZmLARP6c1, a potential RNA-binding protein required for competitive pollen tube growth

**DOI:** 10.1101/2020.11.29.402982

**Authors:** Lian Zhou, Zuzana Vejlupkova, Cedar Warman, John E. Fowler

**Affiliations:** Maize Research Institute, College of Agronomy and Biotechnology, Southwest University, Beibei, Chongqing 400715, China; Department of Botany and Plant Pathology, Oregon State University, Corvallis, OR 97331, USA

**Keywords:** maize, ZmLARP6c1, pollen competition, pollen germination, pollen tube growth

## Abstract

Members of the La-Related Protein family (LARPs) contain a conserved La module, which has been associated with RNA-binding activity. Expression of the maize gene GRMZM2G323499/Zm00001d018613, a member of the LARP family, is highly specific to pollen, based on both transcriptomic and proteomic assays. This suggests a pollen-specific RNA regulatory function for the protein, designated ZmLARP6c1 based on sequence similarity to the LARP6 subfamily in *Arabidopsis*. To test this hypothesis, a *Ds-GFP* transposable element insertion in the *ZmLarp6c1* gene (*tdsgR82C05*) was obtained from the Dooner/Du mutant collection. Sequencing confirmed that the *Ds-GFP* insertion is in an exon, and thus likely interferes with ZmLARP6c1 function. Tracking inheritance of the insertion via its endosperm-expressed GFP indicated that the mutation was associated with reduced transmission from a heterozygous plant when crossed as a male (ranging from 0.5% to 26.5% transmission), but not as a female. Furthermore, this transmission defect was significantly alleviated when less pollen was applied to the silk, reducing competition between mutant and wild-type pollen. Pollen grain diameter measurements and nuclei counts showed no significant differences between wild-type and mutant pollen. However, *in vitro*, mutant pollen tubes were significantly shorter than those from sibling wild-type plants, and also displayed altered germination dynamics. These results are consistent with the idea that ZmLARP6c1 provides an important regulatory function during the highly competitive progamic phase of male gametophyte development following arrival of the pollen grain on the silk. The conditional, competitive nature of the *Zmlarp6c1::Ds* male sterility phenotype (i.e., reduced ability to produce progeny seed) points toward new possibilities for genetic control of parentage in crop production.

## INTRODUCTION

The life cycle of plants alternates between haploid and diploid phases, called the gametophyte and sporophyte, respectively (Walbot and Evans, 2003). Flowering plants have male and female gametophytes, both of which consist of multiple cells and require regulation of gene expression for their function. For the male gametophyte, pollen grain development initiates inside the anther post-meiosis, with the unicellular microspore. In maize, the mature pollen grain consists of a vegetative cell and two sperm cells after two mitoses during its development. When the pollen grain arrives at the maize silk, progamic development commences, as the pollen tube germinates and its growth is directed to the embryo sac for fertilization (Cheung et al., 1995; Hulskamp et al., 1995; Johnson et al., 2019). Proper elongation of the pollen tube is essential for fertilization (Dresselhaus et al., 2016; Hafidh et al., 2016).

Analysis of the male gametophyte transcriptome in multiple plant species, including maize, indicates that there are numerous genes specifically expressed in the male gametophyte, apparently distinct from any expression or function in the sporophyte (Honys and Twell, 2004; Pina et al., 2005; Chettoor et al., 2014; Rutley and Twell, 2015; Warman et al., 2020). Relatively few of these genes have been characterized *in vivo* with respect to their effect on reproductive success, i.e., their contribution to successful pollen tube growth, sperm cell delivery, fertilization and seed set, particularly in monocots (Procissi et al., 2001; Lalanne and Twell, 2002; Kim et al., 2019; Lopes et al., 2019). Notably, mutations in genes active in the male haploid gametophyte are associated with reduced transmission through the male, as loss of important genetic functions can impair the ability to compete with wild-type pollen tubes, therefore reduce reproductive success, i.e., fitness (Arthur et al., 2003; Warman et al., 2020). This reduced transmission can make the generation of mutant homozygotes difficult, and it can also conceivably reduce the recovery or maintenance of such mutations in a population. Notably, although a number of cell biological regulators active during progamic development have been characterized by mutation (Chang et al., 2013; Kaya et al., 2014; Huang et al., 2017), few direct regulators of gene expression enabling pollen tube growth have been genetically defined (Lago et al., 2005; Adamczyk and Fernandez, 2009; Reňák et al., 2012).

In higher plants, post-transcriptional regulation is essential for proper gene expression during floral development and hormone signaling (Fedoroff, 2002; Kramer et al., 2018; Cho et al., 2019; Prall et al., 2019). RNA-binding proteins are associated with post-transcriptional regulation, including pre-mRNA processing, transport, localization, translation and stability of mRNAs (Dreyfuss et al., 2002; Dedow and Bailey-Serres, 2019). In particular, RNA-binding proteins harboring an RNA recognition motif (RRM) at the N-terminus and a glycine-rich region at the C-terminus are thought to play crucial roles in plant growth and stress responses (Ciuzan et al., 2015; Koster et al., 2017).

The La-related proteins (LARP) are a large and diverse superfamily, each member harboring the La module, comprised of two conserved and closely juxtaposed domains that are thought to promote RNA binding: the La motif followed by an RRM motif (Koster et al., 2017; Maraia et al., 2017). The superfamily is further divided into five families: LARP1, LARP3 (encompassing the original La protein), LARP4, LARP6 and LARP7, based on structural features and evolutionary history (Bousquet-Antonelli and Deragon, 2009; Bayfield et al., 2010; Deragon, 2020). Several members of these families have been proven to be RNA-binding proteins. The La protein binds to RNA polymerase III (RNAP III) transcripts at the single-stranded UUU-3’OH (Wolin and Cedervall 2002). LARP7 family members most closely resemble LARP3 but function with a single RNAP III nuclear transcript, 7SK, or telomerase RNA (Krueger et al., 2008; Eichhorn et al., 2016; Eichhorn et al., 2018; Mennie et al., 2018). In animal cells, LARP1 and LARP4 bind to the 5’ and 3’ untranslated regions of mRNAs to regulate translation and degradation (Aoki et al., 2013; Merret et al., 2013b; Lahr et al., 2015). In plants, the roles of the LARP superfamily are only beginning to be characterized. For example, in *Arabidopsis*, the *AtLa1* gene, a true La ortholog in the LARP3 family, is required for completion of embryogenesis, perhaps in part due to its role in promoting translation of the WUSCHEL (WUS) transcription factor, which is required for stem cell homeostasis. At the molecular level, AtLa1 appears to promote translation via interaction with the 5’UTR of the *WUS* mRNA (Fleurdepine et al., 2007; Cui et al., 2015). Members of the *Arabidopsis* LARP1 family have been linked to both leaf senescence and heat-induced mRNA decay (Zhang et al., 2012; Merret et al., 2013a). Most recently, LARP1 has been shown to regulate the translation of the subset of *Arabidopsis* transcripts bearing the 5’-terminal oligopyrimidine stretch (TOP) motif, as a key component of the target of rapamycin (TOR) pathway regulating cellular metabolism (Scarpin et al., 2020).

In contrast, there is no analogous cellular or organismal function yet identified for the LARP6 family. Only one endogenous RNA ligand has been identified: in mammals, LARP6 binds to a secondary structure element in the 5’UTR of type I collagen mRNAs (Cai et al., 2010). The LARP6 interdomain linker between the La motif and the RRM plays a critical role in this RNA binding activity (Martino et al., 2015). Intriguingly, certain members of the plant LARP6 family interact with the major plant poly(A)-binding protein (PAB2), and also show differential RNA-binding activity *in vitro* (Merret et al., 2013b).

Bioinformatics analyses (Merret et al., 2013b) showed that, in the maize genome, there are six genes encoding LARP6 family proteins (ZmLARP6a, ZmLARP6b1-b3 and ZmLARP6c1-c2). Here, we report that one of these genes (GRMZM2G323499/Zm00001d018613), which encodes ZmLARP6c1, is specifically and highly expressed in mature pollen. To begin defining the function of this gene, a *Ds-GFP* transposable element insertion (Li et al., 2013) in its coding sequence was obtained and confirmed. Characterization of this mutation and its phenotypes points toward an important male-specific function for ZmLARP6c1 in the haploid gametophyte during the highly competitive phase of pollen tube germination and growth following pollination.

## MATERIALS AND METHODS

### Plant Materials, Growth Conditions

The *Ds-GFP* allele used in this study (*tdsgR82C05*) was requested from the Maize Genetics Cooperation Stock Center based on its insertion location in the coding sequence of the *larp6c1* gene (acdsinsertions.org). The mutation was backcrossed to a *c1* homozygous, predominantly W22 inbred line at least twice to insure heterozygosity and the presence of a single *Ds-GFP* locus before use in the experiments described. Primers used to PCR-genotype mutant and wild-type alleles in plants from segregating families are listed in Supplementary Table 1. Genetic studies were carried out using standard procedures at the OSU Botany & Plant Pathology Farm in Corvallis, OR.

### Sequence Alignment and Phylogenetic Analysis

The full-length sequences of ZmLARP6 and AtLARP6 family proteins were obtained from MaizeGDB (https://maizegdb.org) and TAIR (http://www.arabidopsis.org/). Multiple amino acid sequence alignment was conducted using MegAlign Software. Phylogenetic analysis was conducted using MEGA6 by neighbor-joining method (Tamura et al., 2013).

### RNA Extraction

Plants for RNA were grown in summer field conditions in Corvallis, OR. Fresh mature pollen was collected upon shedding between 11:00 am and 12:00 pm. 50 mg pollen of each plant was quickly harvested, frozen in liquid nitrogen and stored at −80°C. Total RNA was isolated from mature pollen using Trizol Reagent (Thermo Fisher scientific, Cat no. 15569026) with added PEG (20,000 MW) to 2% (Tattersall et al., 2005) and quantified via Nanodrop (ND-1000).

### Real-time qRT-PCR

First-strand cDNAs were synthesized from total RNA using SuperScript First-strand synthesis system for RT-PCR kit (Invitrogen). Real-time qRT-PCR was performed using SYBR Premix EX TaqTM ⊓ (TAKARA) on a CFX96 Real-Time system (Bio-Rad), according to the manufacturer’s instructions. The amplification program was performed at 95°C for 5 s, 60°C for 30 s and 72°C for 30 s. Triplicate quantitative assays were performed on each cDNA sample. The relative expression was calculated using the formula 2^-Δ(Δcp)^. The housekeeping gene *ZmUbiquitin1* was used as an internal control. All primers used for real-time qRT-PCR are listed in Supplementary Table 1.

### *In Vitro* Pollen Germination

Plants for pollen germination experiments were grown under long day conditions in a greenhouse at OSU or in summer field conditions in Corvallis, OR. Pollen was germinated in pollen germination medium (PGM - 10% sucrose, 0.0005% H3BO3, 10 mM CaCl2, 0.05 mM KH2PO4, 6% PEG 4000) (Schreiber and Dresselhaus, 2003) and imaged at 15 min or 30 min after exposure to media. Germinated, ungerminated and ruptured pollen were quantified from digital images using the FIJI distribution of ImageJ (Schindelin et al., 2012), with a minimum of 125 grains categorized for each datapoint. Pollen tube lengths were measured from homozygous *larp6c1::Ds-GFP* mutants and closely-related wild-type comparator plants. Four replicates were measured with FIJI, with three homozygous mutant and three wild type plants used per replicate, and a minimum of 50 pollen tubes measured for each pollen collection.

### Statistical Analysis

Statistical significance was determined using t-test or lm functions either in SAS (SAS Institute Inc., http://www.sas.com) or with R. Analysis of the categorical response in pollen germination (i.e., the proportion of germinated, ruptured, and ungerminated pollen grains) via logistic regression used the mblogit function (Baseline-Category Logit Models for Categorical and Multinomial Responses) of the mclogit package in R. The categorical dataset was modeled using Genotype + Timepoint + Replicate as factors, with dispersion estimated using the “Afroz” default. R code and the original dataset are available upon request.

### Counting of Fluorescent Kernels

Maize ears were scanned using a custom-built scanner and image projection pipeline (Warman et al., 2020a; Warman et al., 2020b). Green fluorescent (GFP) kernels and non-GFP kernels were counted using the digital image projections and the ‘Cell Counter’ plugin of the FIJI distribution of ImageJ.

### DyeCycle Green Pollen Staining

Fresh pollen was collected from each plant, immediately fixed in ethanol: acetic acid (3:1 ratio) solution, and later rehydrated through an ethanol series for size measurements and staining. To visualize nuclei, fixed pollen grains were stained with DyeCycle Green (Thermo Fisher Scientific, Cat no. V35004) at a final concentration of 1 μM in water, incubated in dark for 30 min and washed with sterile water. Stained pollen was then imaged on a Zeiss Axiovert S100 microscope equipped with HBO 50 Illuminator and Chroma #41017 filter set (470/40 excitation, 495 long-pass dichroic, 525/50 emission) using Qimaging (Retiga ExI) camera.

## RESULTS

### *ZmLarp6c1* Encodes a Pollen-Specific Member of the LARP6 Family

Members of the La-Related Protein family (LARPs) contain a conserved La motif (LAM) and an RNA recognition motif (RRM), which has been associated with RNA-binding activity. GRMZM2G323499/Zm00001d018613 is a member of the LARP family, and is predicted to encode a 505 amino acid protein that harbors conserved La and RRM-like motifs (RRM-L) (**Figure 1A)**. The presence of a conserved glycine-rich C-terminal domain (the LSA: LAM and S1 associated motif) in six members of the maize LARP family is diagnostic for placing them in the LARP6 subgroup (**Figure 1B**). Consistent with earlier analyses, all but one of these also harbors the PABP-interacting motif 2 (PAM2) (**Figure 1B**), which may mediate binding to Poly(A)-Binding Protein (PABP) (Merret et al., 2013). A phylogenetic tree was constructed based on the amino acid sequence alignment of LARP6 subgroups of maize and *Arabidopsis*. Phylogenetic analysis confirms that ZmLARP6c1 has a high degree of sequence similarity with other ZmLARP6s and that the evolutionarily closest *Arabidopsis* protein is AtLARP6c (**Figure 1C**). The closest LARP in maize, designated ZmLARP6c2, is encoded by GRMZM2G411041/Zm00001d032397 on chromosome 1, which is non-syntenic with *ZmLarp6c1* near telomere of the chromosome 7S.

**FIGURE 1.**
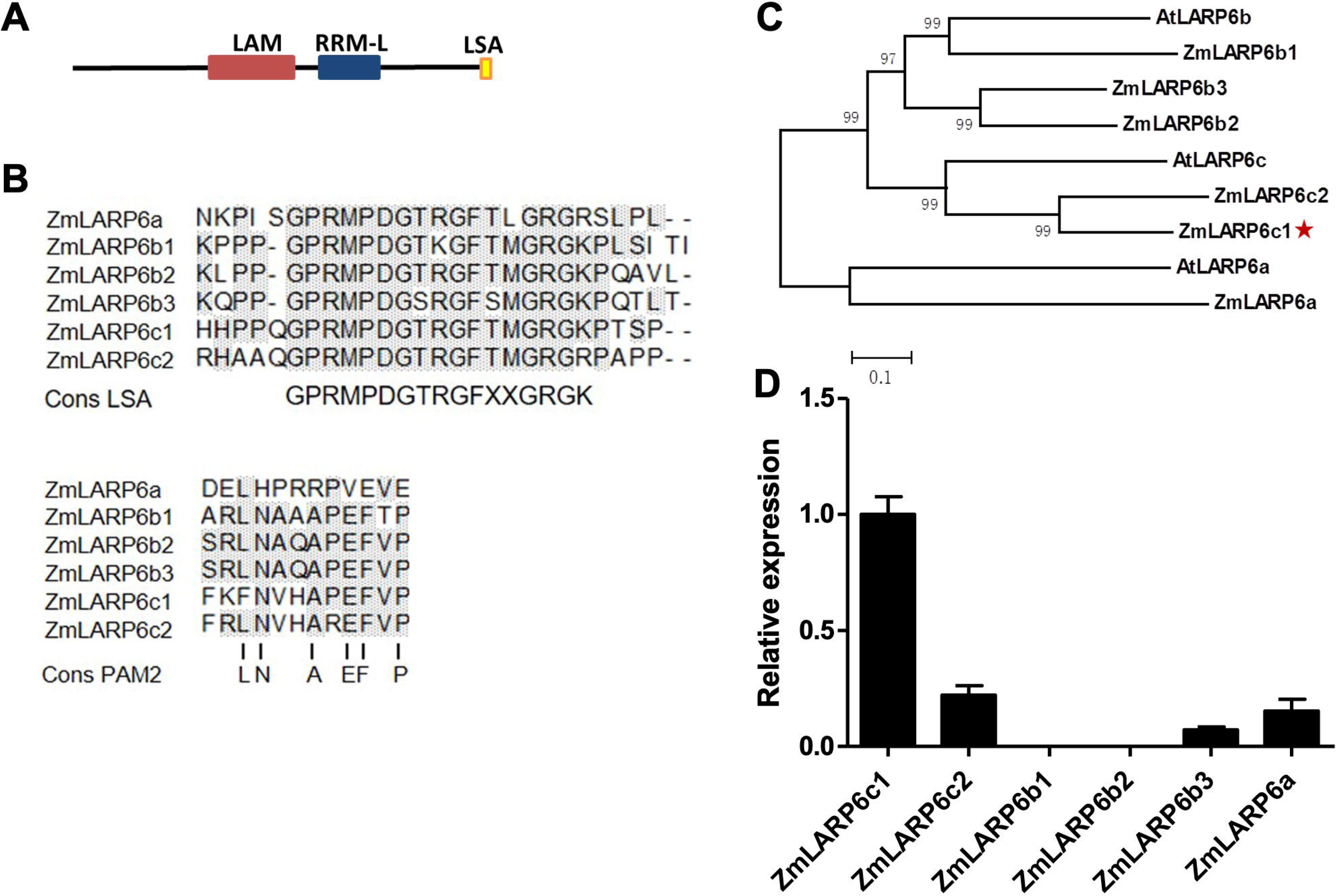
Bioinformatic analysis and expression profiling of the maize LARP6 family members points toward a role for *ZmLarp6c1* in maize pollen. **(A)** Predicted conserved domain architecture of ZmLARP6c1. **(B)** Alignments of maize LARP6 amino acid sequences alongside consensus sequences for the LSA motif and the PAM2 motif. Gray shade, matched residues. **(C)** Phylogenetic tree, using the entire amino acid alignment and the distance method. The bar indicates a mean distance of 0.1 changes per amino acid residue. (**D**) Relative transcript levels of *ZmLARP6* genes in mature pollen of maize. Total RNA was extracted from mature pollen for qRT-PCR. All data are means of three biological replicates with error bars indicating SD.

Expression of *ZmLarp6c1* is highly specific to pollen, based on both transcriptomic and proteomic datasets (**Supplementary Figure 1**; Walley et al 2016, Warman et al 2020). To assess directly expression of all genes encoding members of the LARP6 family in maize pollen, total RNA was isolated from mature pollen collected from inbred line B73. Real-time qRT-PCR results showed that *ZmLarp6c1* had the highest transcriptional abundance of all six *ZmLarp6* family members in mature pollen (**Figure 1D**). Together, these data suggest a pollen-specific RNA regulatory function for Zm LARP6c1.

### A *ZmLarp6c1* Mutation is Associated with Reduced Transmission when Crossed as a Male

To investigate the function of *ZmLARP6c1* during pollen development, a line carrying a *Ds-GFP* transposable element insertion in *ZmLARP6c1* (*Zmlarp6c1::Ds-GFP*) was obtained from the Dooner/Du population (Li et al., 2013). Sequencing confirmed that the *tdsgR82C05* insertion is in the fifth exon (**Figure 2A**) in the RRM-L domain coding region, and thus likely interferes with *ZmLARP6c1* function. PCR genotyping primers were designed to confirm the co-segregation with the GFP marker, and identify the genotype (**Figure 2B**).

**FIGURE 2.**
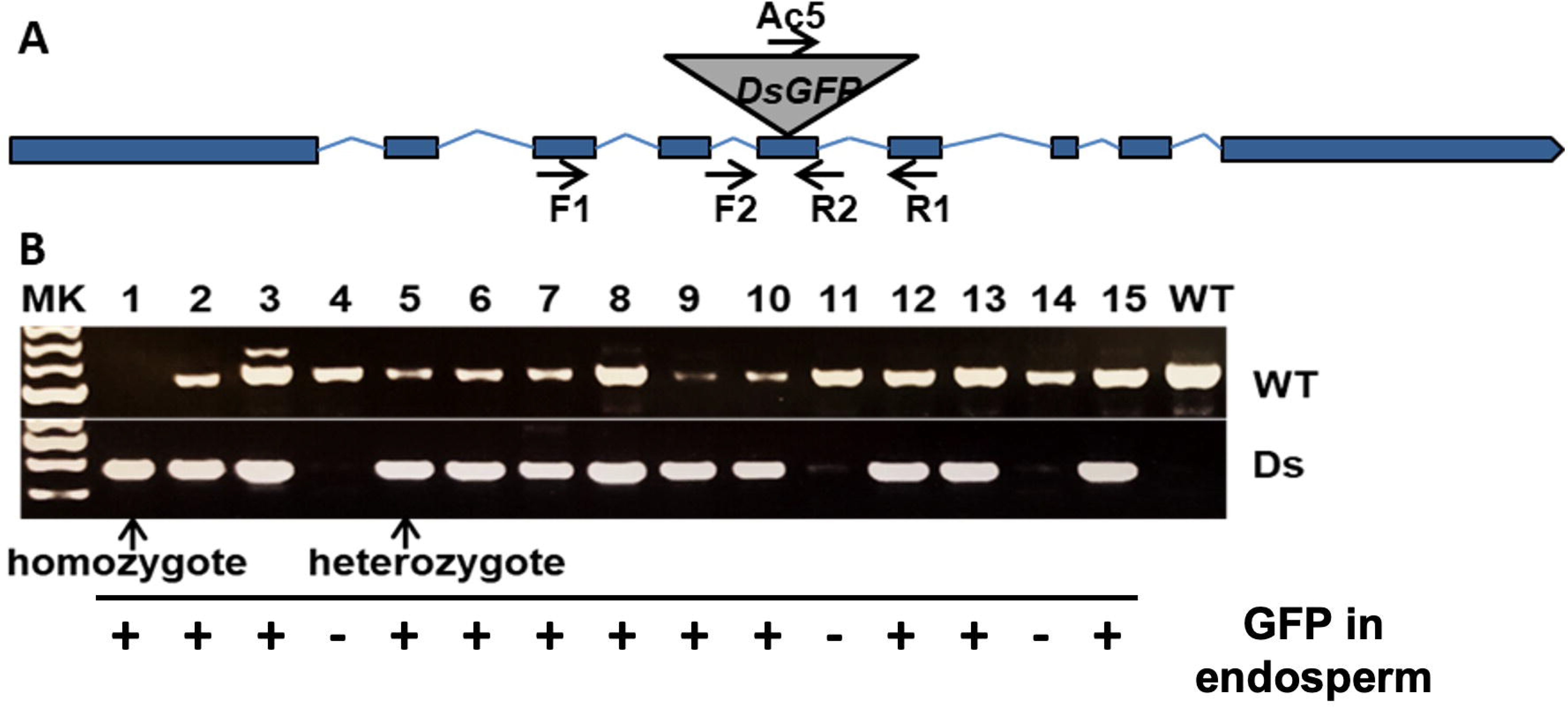
A *Ds-GFP* transposable element insertion line provides a tool to investigate *Zmlarp6c1* function. **(A)** Sequencing confirmed that the insertion was in the 5^th^ exon; thus, the mutation, likely a knockout, could be easily tracked genetically by observing GFP fluorescence in the endosperm. **(B)** PCR genotyping of progeny (using primers F1, R1 and Ac5) from a selfpollinated heterozygote confirms co-segregation of the insertion with the GFP marker, and helps clarify genotype (homozygote vs. heterozygote).

Neither heterozygous nor homozygous *Zmlarp6c1::Ds-GFP* plants showed any obvious differences in vegetative and floral development. As an initial assessment of effects on gametophytic development and function, *Zmlarp6c1::Ds-GFP* heterozygous mutant plants were reciprocally crossed with wild-type plants in the field. Inheritance of the insertion was tracked via its endosperm-expressed GFP, with the mutant and wild-type kernels quantified via annotation of 360° ear projections (**Figure 3**). When crossed through the male, GFP transmission ranged from 0.5% to 26.5%, showing significantly reduced transmission in every outcross in two field seasons (χ^2^ test, p-values <10^-6^), averaging 12.4% transmission across nine ears (**Table 1**). In contrast, female GFP transmission ranged from 47.2% to 56.7%, averaging 51.4%, with only a single cross significantly different from expected Mendelian inheritance (56.7%, p = 0.008) (**Table 1**).

**FIGURE 3.**
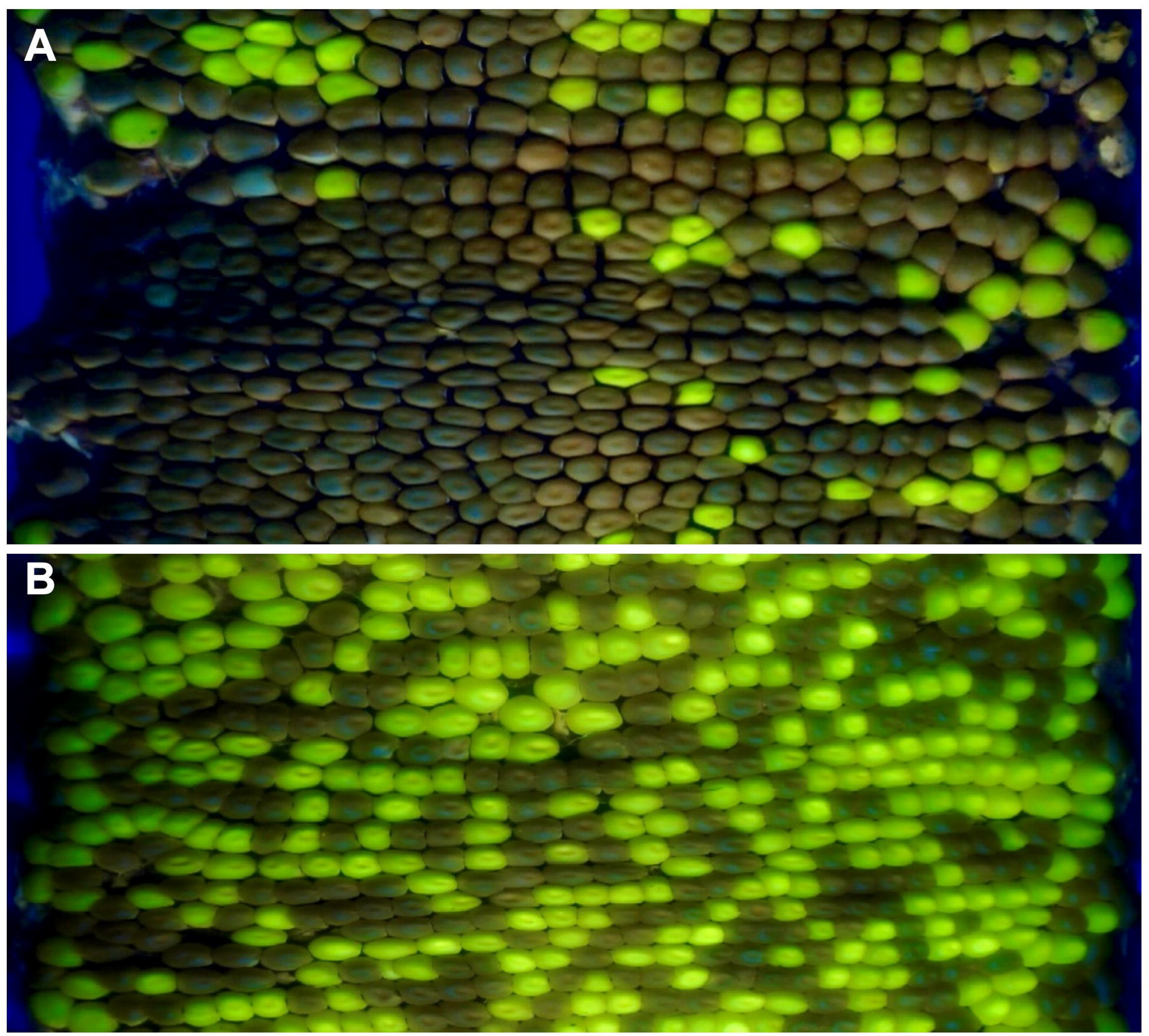
Maize ear projections demonstrate a strong male-specific transmission defect associated with the GFP-expressing *Zmlarp6c1::DsgR82CO5* allele. **(A)** *Zmlarp6c1::Ds/+* crossed as a male (52 GFP kernels/420 total kernels = 12.4% transmission). **(B)** The same *Zmlarp6c1::Ds/+* plant crossed as a female (246 GFP kernels/479 total kernels = 51.4% transmission).

**TABLE 1.** Transmission of *larp6c1::DsGFP* in reciprocal outcrosses.

| *larp6c1::DsGFP*/+<br>plant/season | Direction of outcross<br>to wild-type | # GFP<br>kernels | # wt<br>kernels | % GFP | p-value ( $\chi^2$ ) |
| --- | --- | --- | --- | --- | --- |
| 1 / 2018 | Male | 30 | 83 | 26.5 | <10<sup>-6</sup> |
|  | Female | 124 | 98 | 55.9 | 0.081 |
| 2 / 2018 | Male | 32 | 160 | 16.7 | <10<sup>-6</sup> |
|  | Female | 187 | 209 | 47.2 | 0.269 |
| 3 / 2018 | Male | 19 | 257 | 6.9 | <10<sup>-6</sup> |
|  | Female | 238 | 228 | 51.1 | 0.643 |
| 4 / 2018 | Male | 63 | 285 | 18.1 | <10<sup>-6</sup> |
|  | Female | 184 | 193 | 48.8 | 0.643 |
| 1 / 2019 | Male | 38 | 119 | 24.2 | <10<sup>-6</sup> |
|  | Female | 221 | 211 | 51.2 | 0.630 |
| 2 / 2019 | Male | 12 | 439 | 2.7 | <10<sup>-6</sup> |
|  | Female | 224 | 171 | 56.7 | 0.008 |
| 3 / 2019 | Male | 52 | 368 | 12.4 | <10<sup>-6</sup> |
|  | Female | 246 | 233 | 51.4 | 0.553 |
| 4 / 2019 | Male | 9 | 256 | 3.4 | <10<sup>-6</sup> |
|  | Female | 221 | 210 | 51.3 | 0.596 |
| 5 / 2019 | Male | 1 | 187 | 0.5 | <10<sup>-6</sup> |
|  | Female | 148 | 154 | 49.0 | 0.730 |

To create derivative excision alleles to confirm that the insertion is causal for the male-specific transmission defect, *Zmlarp6c1::Ds-GFP* was crossed into an *Ac-im* active line (Conrad and Brutnell, 2005). Progeny were screened by PCR for excision, and 6 putative revertant alleles were identified. Two of these alleles were heritable and confirmed by sequencing (**Figure 4A, B**). As expected, there was an 8 bp duplication in the original *Zmlarp6c1::Ds-GFP* insertion allele compared to wild-type. *Ds* excision generated distinct nucleotide footprints in the two derivatives, with one footprint (*dX549B2*) adding six base pairs relative to the wild-type sequence and the other (*dX550A8*) adding eight base pairs (**Figure 4B**). Given an insertion in the coding region, the eight-bp insertion would be predicted to generate a frameshift, and thus a likely non-functional LARP6c1 protein, whereas the six-bp footprint would insert two amino acids but retain the frame (**Supplementary Figure 2**), and thus potentially restore protein function. To test for altered functionality associated with these variant sequences, plants heterozygous for each of the two alleles were outcrossed as males with a heavy pollen load, and transmission was assessed via PCR genotyping. As expected, the eight-bp footprint *dX550A8* allele retained a strongly reduced transmission rate (12.5%, **Figure 4C**). However, the six-bp footprint *dX549B2* allele recovered a wild-type transmission rate (54.2%, **Figure 4D**), indicating reversion of gene function. These data confirm that the interruption of the *Zmlarp6c1* coding sequence by the original *tdsgR82C05* insertion was causal for the transmission defect, rather than an unknown, linked mutation.

**FIGURE 4.**
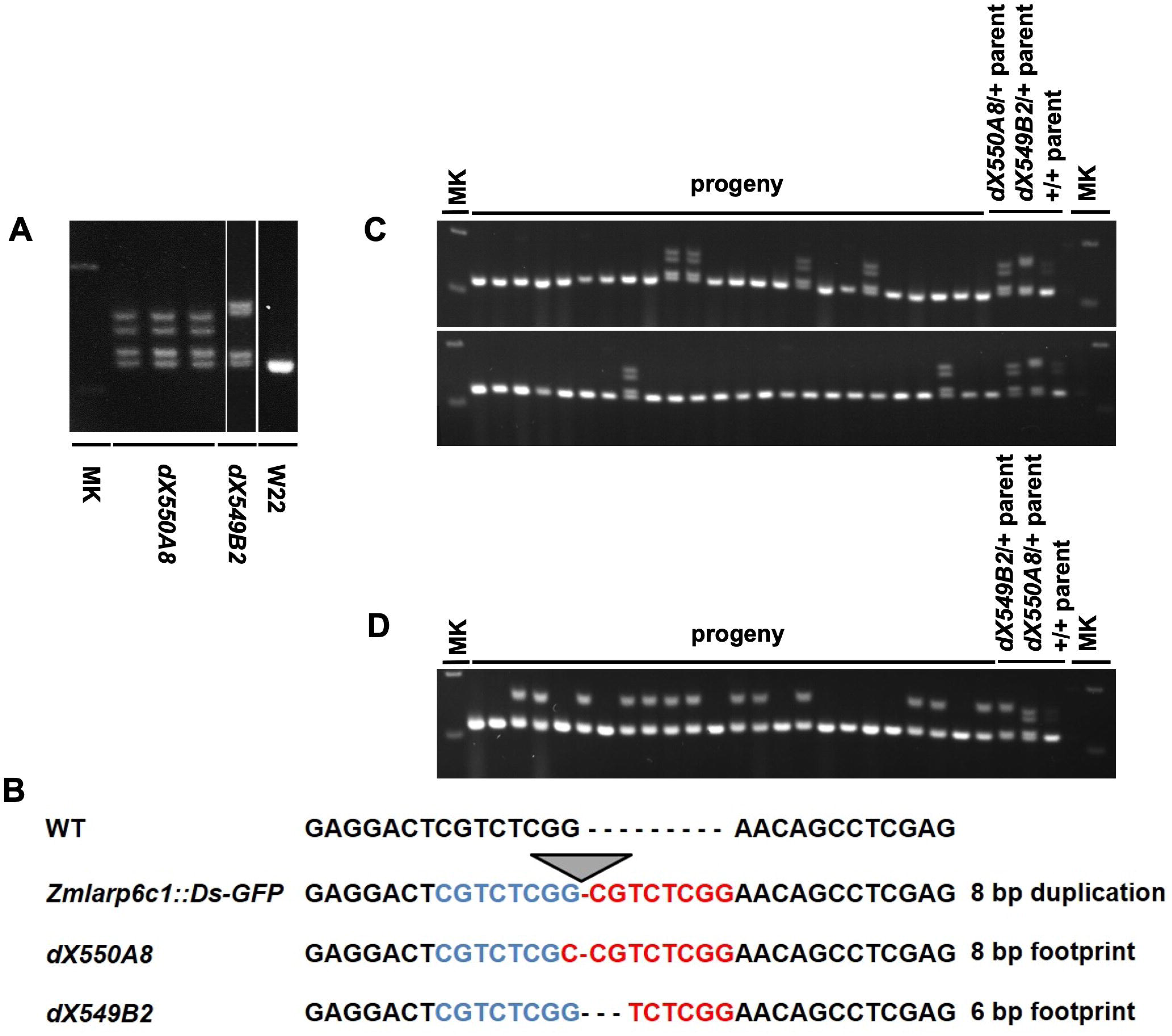
*Ds* excision alleles derived from *Zmlarp6c1::DsgR82C05* validate that the male specific transmission defect is due to loss of ZmLARP6c1 function. **(A)** PCR genotyping using primers F2 and R2 (Figure 2) documents isolation of two derivative alleles (*dX550A8* and *dX549B2*) due to excision of the *Ds-GFP* element from *Zmlarp6c1::DsgR82C05*. W22: comparator wild-type inbred; MK: DNA size marker. **(B)** Sequencing of the derivative alleles shows that *dX550A8* and *dX549B2* are associated with an eight-bp and six-bp footprints, respectively, in the *Zmlarp6c1* coding sequence. **(C)** *Zmlarp6c1-dX550A8* is associated with a male-specific transmission defect, with 6 out of 48 progeny from a heterozygous male outcross inheriting the footprint (12.5%). **(D)** *Zmlarp6c1-dX549B2* segregates at a Mendelian ratio when transmitted through the male (13:11 from a heterozygous outcross).

### *ZmLARP6c1* Contributes to Pollen Competitive Ability *in vivo* and to Robust Pollen Tube Growth *in vitro*

Notable in the transmission tests was the observation that the male transmission rate was associated with high variance (standard deviation = 9.6%), relative to female transmission (standard deviation = 3.1%). This raised the possibility that *Zmlarp6c1::Ds* pollen is particularly sensitive to an environmental factor that was variable across these ears. One such factor that can be easily tested is the level of competition – i.e., the number of wild-type pollen grains competing with mutant pollen grains on each silk. In four independent experiments, a single pollen collection from a mutant heterozygote was used to heavily pollinate silks of one ear (HVY), and to sparsely pollinate three other ears with limiting pollen (SPS) to reduce competition. In three of the four experiments, sparse pollination significantly relieved the GFP transmission defect, relative to the comparator heavy pollination, with transmission rates from 37.7% to 53.3% (**Table 2**). Because these experiments are inherently imprecise, we sought to find a measure to assess pollen load for each pollination. The total number of seeds per ear provides such an assessment, indirectly, given that seed production is limited to ovules successfully pollinated and fertilized. Analysis of all ears in the dataset individually showed a significant correlation between GFP transmission rate and the total number of seeds on an ear (**Figure 5**) (p= 0.0003). This is consistent with a strong effect of the level of competition with wild-type pollen on the ability of *Zmlarp6c1::Ds* pollen to successfully fertilize and generate a seed. These data argue that *Zmlarp6c1* is a crucial contributor to pollen competitive ability *in vivo*, and suggest that its function is manifested post-pollination.

**FIGURE 5.**
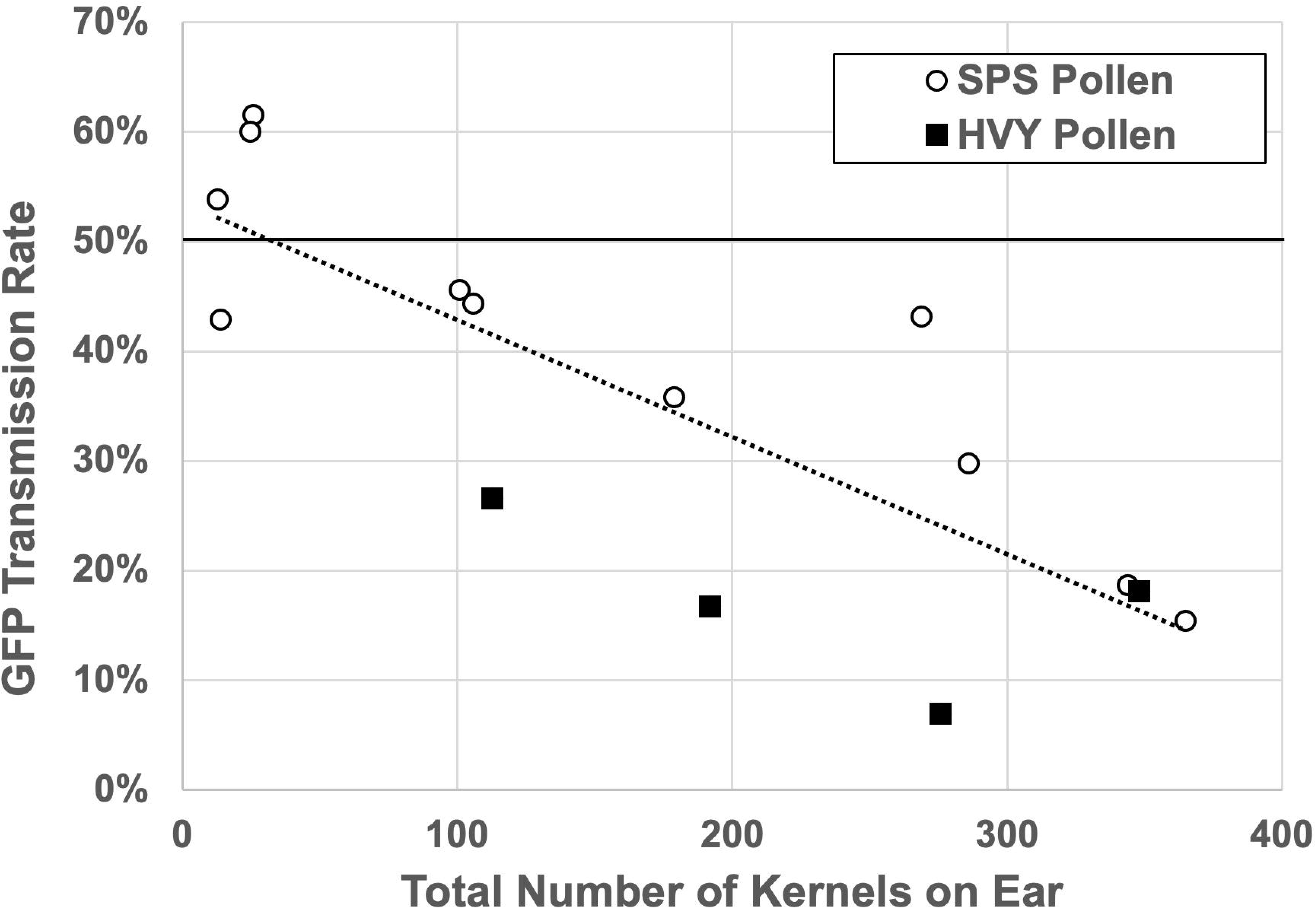
Transmission of *Zmlarp6c1::DsgR82C05* mutation is sensitive to pollen load. Outcomes of each separate ear with more than 10 kernels in the experiments from Table 2 are plotted. Pollination with heavy, near-saturating (HVY) pollen is associated with reduced transmission of *Zmlarp6c1::Ds*, relative to sparse (SPS) pollen (t-test, p = 0.002601). Moreover, linear regression (dotted line) indicates a significant correlation between total kernels per ear and transmission rate (p = 0.0003, R^2^ = 0.647).

**TABLE 2.**
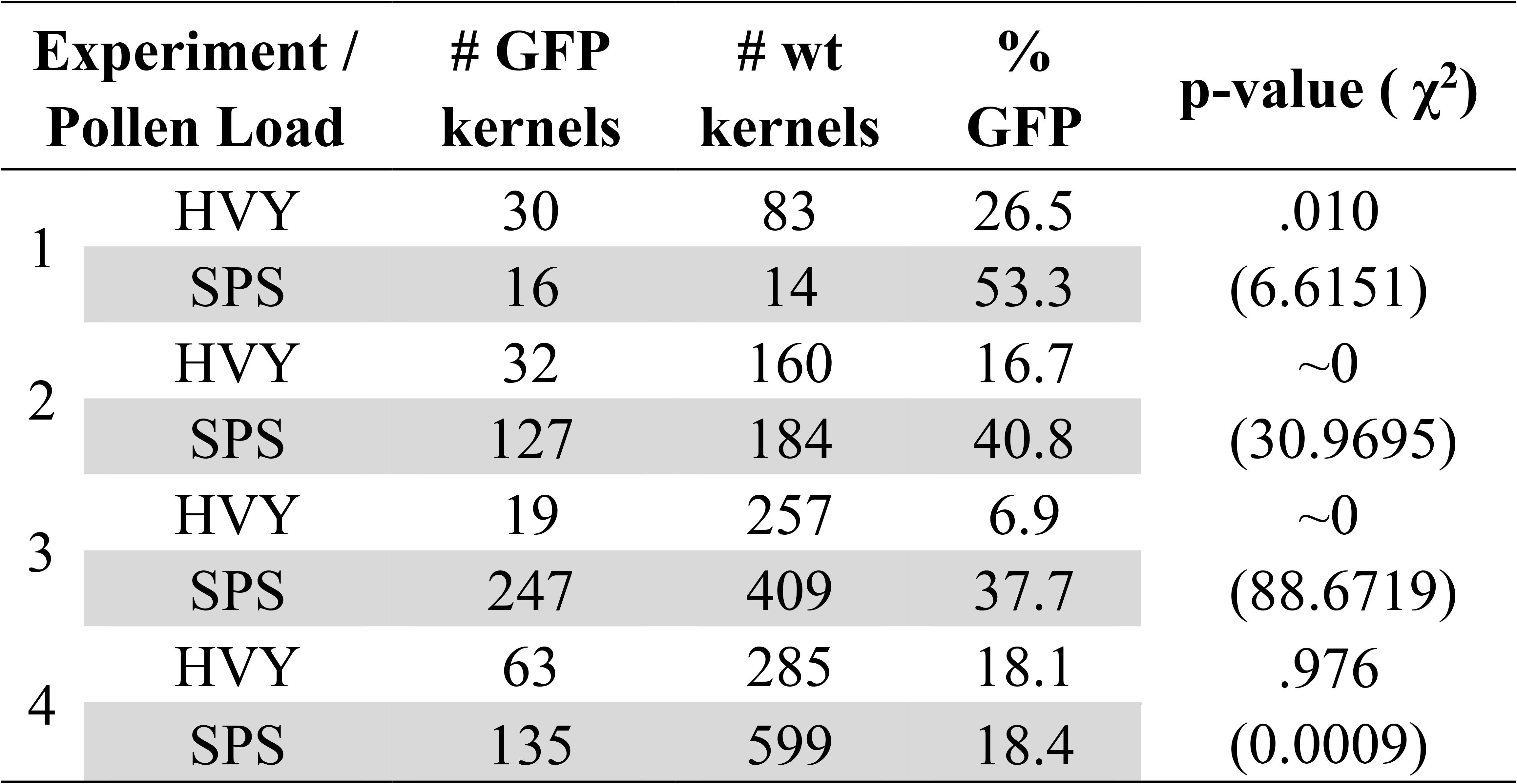
Transmission of *larp6c1::DsGFP* through the male when pollen load is varied.

One possibility to explain the competitive defect would be slowed pollen tube germination, and/or a reduced growth rate for pollen tubes. To assess these possibilities, we characterized pollen tube germination and growth *in vitro* (**Figure 6**). *In vitro* pollen was scored at 15 min and 30 min after exposure to pollen growth medium (PGM), with germinated, ungerminated and ruptured pollen grains counted. (Ruptured pollen grains have ceased growing after releasing cytoplasm into the media, apparently due to loss of cell wall integrity at the growing pollen tube tip.) In initial experiments, pollen germination rate from the heterozygous *Zmlarp6c1::Ds-GFP* plants (30.3% and 46.4%) was significantly lower than that from a sibling wild-homozygote (59.3% and 71.7%) at 15 min and 30 min, respectively (**Supplementary Figure 3**). This lower germination rate was accompanied by increased ungerminated pollen from the mutant, rather than significant changes in the percentage of ruptured grains (**Supplementary Figure 3)**.

**FIGURE 6.**
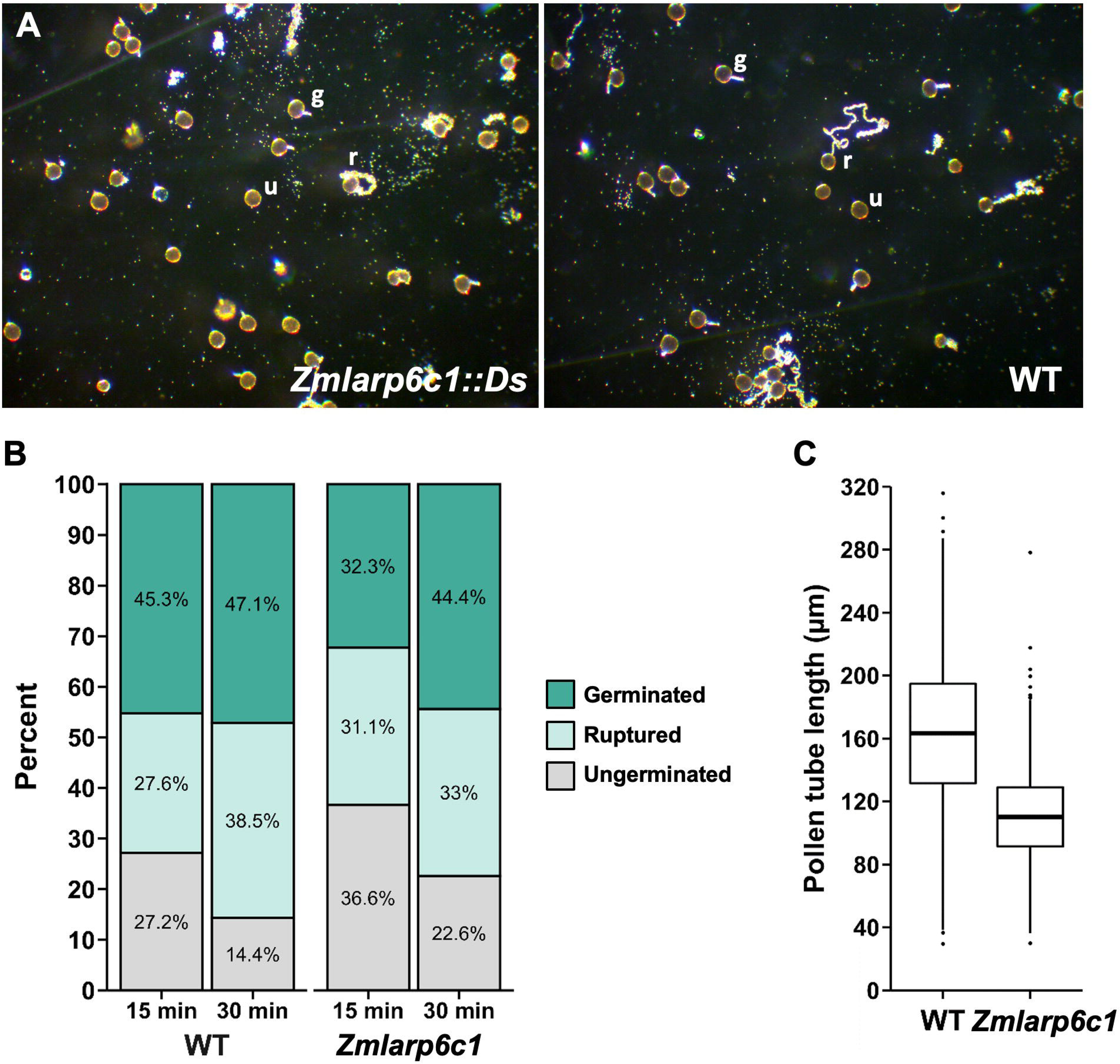
*In vitro*, pollen from *Zmlarp6c1::Ds* plants is associated with shorter pollen tubes and altered germination dynamics. **(A)** Representative fields of pollen germinated *in vitro*, at 15 minutes post-exposure to liquid media. Pollen was collected from either a homozygous *Zmlarp6c1::Ds* plant, or a comparator wild-type (WT) plant. Fields were scored blindly (four biological replicates of three plants of each genotype, at least 125 pollen grains from each plant), categorizing pollen grains as germinated, ruptured, or ungerminated. **(B)** Combined data from four biological replicates, at 15 and 30 minutes after plating in PGM. Modeling the categorical response using a multinomial baseline-category logit model indicates genotype is a significant predictor (p = 0.0047), as is timepoint (p= 1.02e-05). **(C)** Mutant pollen tube lengths at 30 minutes were significantly shorter than wild-type in all replications (Welch’s t-test, all p-values < 10^-15^). Averaged across all four replicates, mutant pollen tubes were 31.5% shorter than wild-type sibling tubes.

A followup experiment with pollen from homozygous field-grown plants confirmed these initial observations, indicating altered germination dynamics associated with loss of *Zmlarp6c1* function (**Figure 6A, B**). Analysis of four replicates comparing wild-type and *Zmlarp6c1::Ds* pollen grown *in vitro* shows the expected significant effect of time on germination, with significant decrease in the percentage of ungerminated pollen from 15 to 30 minutes in both genotypes (p = 1.02E-05), along with a concomitant increase in germinated and ruptured pollen (**Figure 6B**). Genotype was also a significant predictor of germination response (p = 0.0047), confirming that *Zmlarp6c1::Ds* pollen is associated with a higher proportion of ungerminated pollen grains. Notably, although the proportion of ungerminated *Zmlarp6c1::Ds* pollen was higher than wild-type at both time points, by 30 minutes the proportional response in the mutant was similar to that of wild-type pollen at 15 minutes (**Figure 6B,** and individual replicates in **Supplementary Figure 4**). These data are consistent with the *Zmlarp6c1::Ds* defect causing a delay in pollen tube germination, rather than an absolute inhibition of germination. Finally, pollen tube lengths were measured at the 30 minute timepoint for both *Zmlarp6c1::Ds-GFP* mutant and wild-type populations. Consistent with the competitive defect detected *in vivo*, mutant pollen was associated with significantly shorter pollen tubes than wild-type, with an average 31.5% tube length reduction associated with loss of *Zmlarp6c1* function (**Figure 6C, Supplementary Figure 5**).

As an initial assessment as to whether the *Zmlarp6c1::Ds-GFP* mutant pollen was associated with defects prior to the progamic phase (i.e., aberrant pollen grain development, potentially leading to reduced competitiveness following pollination), we measured pollen grain diameter and determined nuclei number in pollen fixed upon anthesis. No significant differences were found in pollen grain diameters comparing sibling wild-type and *Zmlarp6c1::Ds-GFP* heterozygotes, nor did Dyecyle Green staining of DNA identify significant differences in the number of nuclei between these pollen populations (**Figure 7**). Collectively, these findings suggest that the important role played by *ZmLARP6c1* in male gametophyte function is primarily manifested in the post-pollination, progamic phase of gametophytic development, rather than in the development of the pollen grain itself.

**FIGURE 7.**
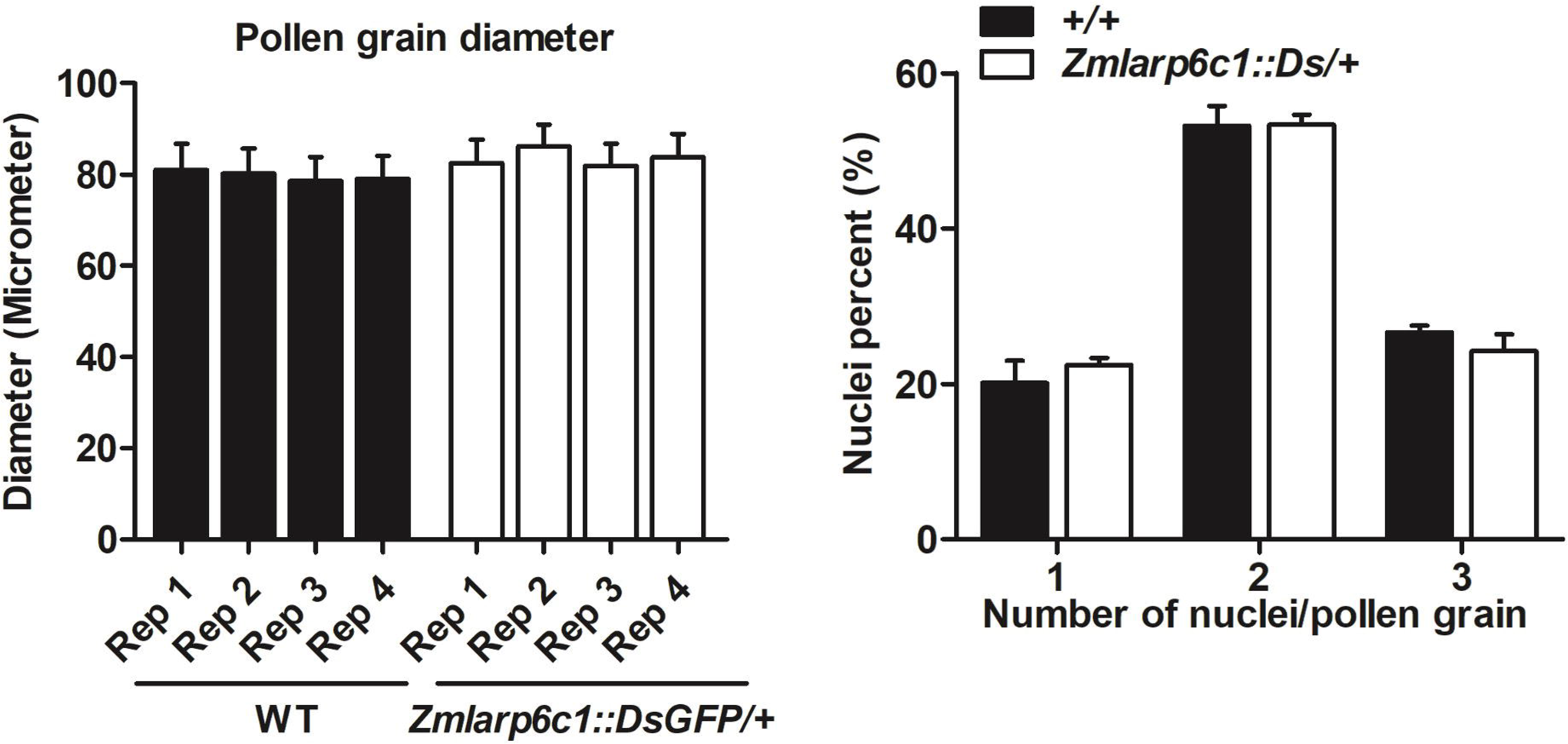
Pollen grain diameter and nuclei count showed no differences between wild-type and mutant pollen. **(A)** At least 150 pollen grains were measured per replicate. **(B)** DyeCycle green fluorescent dye was used to stain DNA in fixed pollen, and distinct nuclei visible in each grain were counted blindly. Although maize pollen grains are expected to have three nuclei at maturity (vegetative cell plus two sperm cells), not every nucleus is discernible in every grain, due to the large size of the pollen grain and presumed variation in staining efficiency. All data are means of four biological replicates with error bars indicating SD.

## DISCUSSION

Flowering plants have a complex sexual reproduction system, which includes the short but critical gametophytic generation, involving pollination followed by double fertilization, triggering development of the progeny seed. Notably, gametophytes express a significant subset of genes in plant genomes, which although reduced relative to the sporophyte (Honys and Twell, 2003; Chettoor et al., 2014; Rutley and Twell, 2015), enable developmental programs and environmental responses. Given that male gametophytic gene expression occurs after the genetic recombination of meiosis, and in organisms that are independent and have a free-living stage, there is considerable potential for expression of phenotypes, upon which evolutionary forces can act (Reese and Williams, 2019). In maize, with its prodigious production of large populations of pollen, lack of a biotic vector, and the extended stigma surface and transmitting tract of the female silk, there appears to be ample opportunity for gametophytic competition in the progamic phase of development (i.e., during pollen tube germination and growth). Assessing transmission rates can give an initial measure of the magnitude of a mutation’s effect on gametophytic fitness (and presumably, biological function), and the same measure can also be used to determine whether competition itself influences fitness upon manipulation of pollen load (Arthur et al., 2003; Cole et al., 2005). The seed-expressed GFP phenotype linked to the *Ds* mutagenic element in the publicly-available *Ds-GFP* population (Li et al., 2013), not only facilitates the initial determination of mutant transmission rate (Warman et al., 2020a), but also broadens the feasibility and precision of analytical approaches for assessing the influence of pollen competition on defects like that associated with *Zmlarp6c1::Ds-GFP* (**Figure 5**).

The biological processes that enable gametophytic function (and the genes that underlie them) are regulated developmentally, but the associated regulatory mechanisms remain unclear. Interestingly, evidence in multiple species indicates that a significant component enabling the transition from quiescent pollen grain to actively growing pollen tube is post-transcriptional control (Hafidh et al., 2016). In pollen of tobacco and *Arabidopsis*, cytoplasmic complexes are likely mechanistically important for storage of transcripts generated earlier in development, and eventual translation during pollen tube growth (Honys et al., 2009; Scarpin et al., 2017). Although such complexes have not been observed in maize pollen, the rapid germination response *in vitro* (pollen tube growth apparent in less than 15 minutes) (**Figure 6**) is consistent with a transition that does not require *de novo* transcription. The detection of significant proteomic differences in germinated maize pollen at 30 minutes after exposed to media is also suggestive that control of translation plays a role in the regulation of pollen tube growth in maize (Walley et al 2016). The protein homology, expression pattern and mutant phenotypes documented in this study raise the possibility that *ZmLarp6c1* is a component of a translational control and/or RNA storage regulatory mechanism.

Few putative RNA-binding proteins have been characterized in detail in mature pollen in any species (Park et al., 2006), and to our knowledge, none have been phenotypically associated with aberrant progamic development. However, a number of conserved sequence elements in transcript UTRs with potential or proven roles in influencing mRNA translation or stability have been identified (Hulzink et al., 2003), most notably in the tobacco *ntp303* transcript (Hulzink et al., 2002). These are potential interaction sites for RNA-binding proteins, possibly including those of the LARP family (e.g., as observed for AtLARP1 and the TOP motif - Scarpin et al., 2020). The severe transmission defect for the *Zmlarp6c1::Ds* mutation indicates that the encoded protein is critical for wild-type pollen function *in vivo*. Moreover, the cellular phenotypes detected via assessment of *Zmlarp6c1::Ds* pollen germination and growth *in vitro* indicate that the encoded protein plays a role in maize pollen post-germination, causing delayed germination dynamics and shorter pollen tubes when mutant. This is in contrast to the lack of a significant effect on two cellular parameters that indicative of effects earlier in development, pollen grain diameter and number of nuclei. Comparing across individual replicates, *Zmlarp6c1::Ds* pollen also appeared to show a higher variance in germination response, relative to the wild-type, raising the possibility that loss of *Zmlarp6c1* function is associated with a more general loss of cellular robustness during pollen tube growth *in vitro* (**Supplementary Figure 4**). Testing these possibilities will require investigating the putative RNA targets and putative binding activity of ZmLARP6c1 in mature pollen and growing pollen tubes.

Genetically-encoded male sterility is of interest for plant breeding, crop production, and protection of patented varieties, for example, in the production of hybrid seeds. Currently, female lines with cytoplasmic male sterility (cms) or genic male-sterility based on expression in the sporophyte (gms) are the preferred biological approaches for insuring only a single male parent during hybrid seed production, and obviate the need for mechanical detasseling (Wan et al., 2019). The observation that a single mutation such as *Zmlarp6c1::Ds* can cause a severe male-specific competitive, rather than absolute, defect raises the possibility of an alternative approach for controlling seed production via genic male sterility based on gametophytic expression (which we are designating ‘gms-g’). The conditional nature of the gametophytic *Zmlarp6c1::Ds* defect - i.e., its ability to generate progeny is hindered only in the presence of wild-type pollen - could be exploited to facilitate not only the generation of hybrid seed, but also seed for the parent female inbred line. This contrasts with sporophytic-based male-sterility of typical gms lines, in which production of any viable pollen is inhibited, complicating the large-scale production of these lines via self-pollination (Wan et al., 2019). In this scenario, a gms-g line that is homozygous for a competition-defective male gametophyte can easily generate seed at scale via open self-pollination in plots isolated from wild-type pollen, eliminating competition. However, for hybrid seed production, the competitive male defect in the female gms-g parent would inhibit productive self-pollination in the presence of wild-type pollen from the hybrid partner male inbred. Similarly, such a genetically-encoded competitive male gametophytic defect could be utilized for discouraging accidental transgene flow or loss of other patented alleles into non-target seed stock. It remains to be seen whether, in an open pollinated field that also provides a robust wild-type pollen source, a defective *Zmlarp6c1* alone is a strong enough barrier to insure effective male sterility.

## Supporting information

Primer Sequences

## AUTHOR CONTRIBUTIONS

JF, CW and LZ conceived and designed the experiments. LZ, ZV, and CW conducted the experiments and analyzed the data. LZ, CW and JF discussed and wrote the manuscript. All authors edited, read and approved the manuscript.

## ACKNOWLEDGMENTS

The authors thank the Dooner/Du collection for the *Ds-GFP* transposable element insertion mutant *tdsgR82C05*. The authors would also like to acknowledge statistical guidance from D. Jiang, OSU Statistics. Finally, thanks to S. Leiboff and other members of the OSU Botany & Plant Pathology PF1 group for their helpful questions and comments.

## SUPPLEMENTARY MATERIALS

**Supplementary Table 1 | Primers used in this study.**

**Supplementary Figure 1 | *Zmlarp6c1* expression is enriched specifically in mature pollen. (A)** Expression profiling data from the Walley et al. (2016) maize developmental atlas, for GRMZM2G323499/Zm00001d018613. RNA-seq data for all 23 different tissue types assessed (blue), with corresponding proteomic (orange) and phosphoproteomic (gray) profiling data for the same tissues. Two additional, relevant samples were assessed for proteomic data: B73 (inbred line) Germinated Pollen and W23 (inbred line) Mature Pollen. Although *Zmlarp6c1* transcript is detected in mature leaf, the translated protein is only detected in pollen samples. FPKM, fragments per kilobase of exon model per million reads mapped; dNSAF, distributed normalized spectral abundance factor. **(B)** Focused transcriptome profiling of maize male gametophyte development (Warman et al. 2020a) indicates that the *Zmlarp6c1* transcript (Zm00001d018613_T004) is highest in mature pollen, but largely excluded from the sperm cells isolated from mature pollen. This suggests the transcript is specifically enriched in the pollen vegetative cell, which drives pollen tube germination and growth.

**Supplementary Figure 2 | Alignments of WT, *dX550A8* and *dX549B2* amino acid sequences.** Red shade, La motif; blue shade, RRM-L; black dot shade, altered residues in derivative alleles; orange oval, two amino acids added by the *dX549B2* six bp footprint at the insertion site.

**Supplementary Figure 3 | Pollen germination rate from a heterozygous *Zmlarp6c1::Ds-GFP* plant was lower than from wild-type plant**. Pollen from sibling plants were germinated in pollen growth medium for 15 min or 30 min. Germinated, ungerminated and ruptured pollen percentages were counted (t-test, p-values < 0.01 for categories marked by an asterisk).

**Supplementary Figure 4 | *Zmlarp6c1::Ds-GFP* and wild-type pollen germination, separated by replicate.** Variance across each replicate is apparent. However, all show a trend toward a smaller percentage of ungerminated pollen grains at 30 min than at 15 min, in both genotypes; and *Zmlarp6c1* pollen associated with a higher fraction of ungerminated pollen than wild type. Modeling the categorical response using a multinomial baseline-category logit model indicates Replicate 2 is a significant factor predictive of a high proportion of Ruptured grains (p = 0.02059), whereas Replicate 4 is a significant factor predictive of a low proportion of Ungerminated grains (p = 0.0134).

**Supplementary Figure 5 | Pollen from homozygous *Zmlarp6c1::Ds-GFP* plants is associated with shorter pollen tubes.** Pollen from homozygous *Zmlarp6c1::Ds-GFP* and comparator wild-type plants was germinated in pollen growth medium for 30 minutes, imaged, and pollen tube lengths from four replicates were measured. Mutant pollen tubes were significantly shorter than wild type in all replicates (Welch’s t-test, all p-values < 10^-15^).

