## Supplementary Figures for "A maize male gametophyte-specific gene encodes ZmLARP6c1, a potential RNA-binding protein required for competitive pollen tube growth"

### Supplementary Figure 1

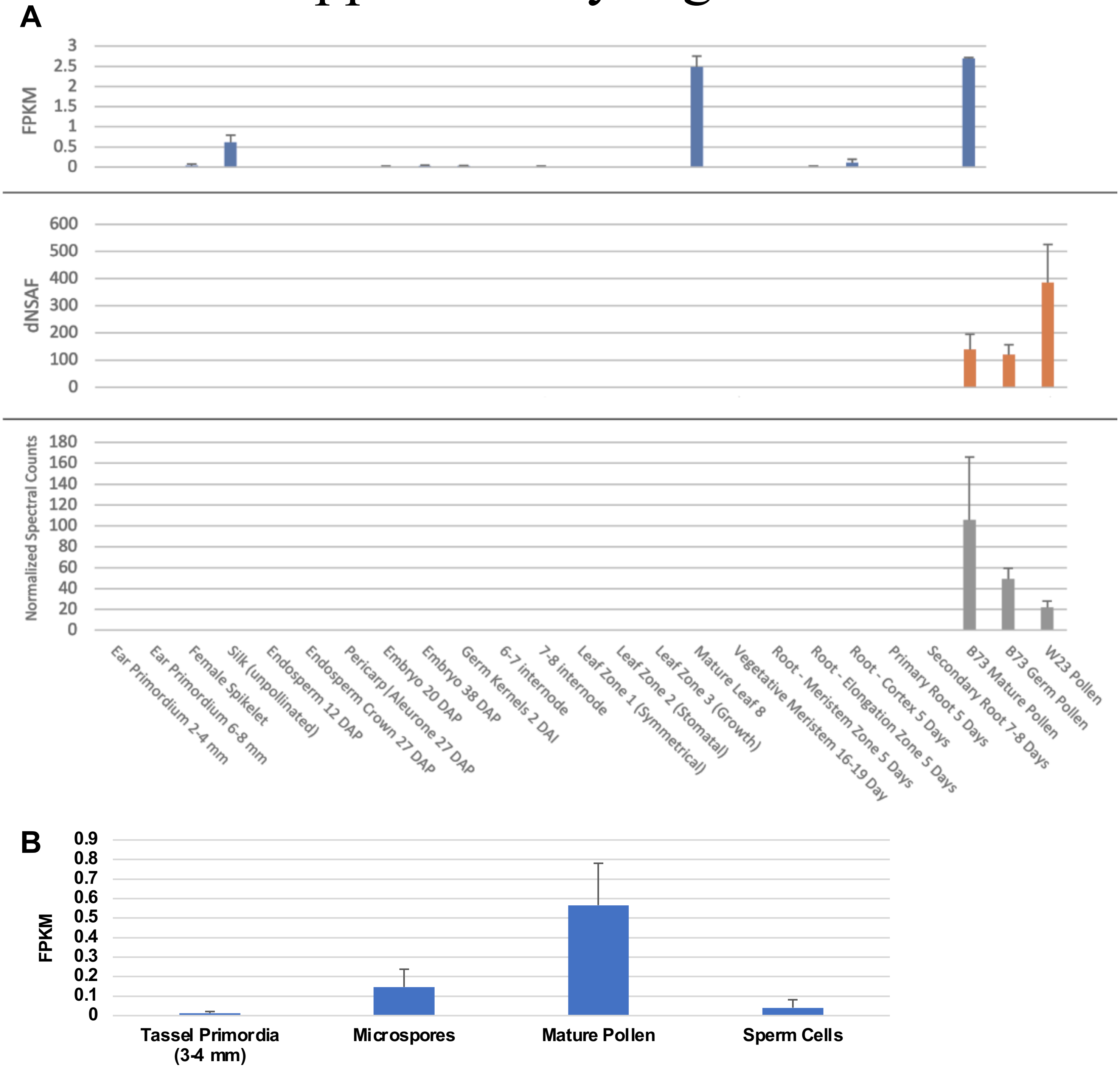


**Supplementary Figure 1 | *Zmlarp6c1* expression is enriched specifically in mature pollen. (A)** Expression profiling data from the Walley et al. (2016) maize developmental atlas, for GRMZM2G323499/Zm00001d018613. RNA-seq data for all 23 different tissue types assessed (blue), with corresponding proteomic (orange) and phosphoproteomic (gray) profiling data for the same tissues. Two additional, relevant samples were assessed for proteomic data: B73 (inbred line) Germinated Pollen and W23 (inbred line) Mature Pollen. Although *Zmlarp6c1* transcript is detected in mature leaf, the translated protein is only detected in pollen samples. FPKM, fragments per kilobase of exon model per million reads mapped; dNSAF, distributed normalized spectral abundance factor. **(B)** Focused transcriptome profiling of maize male gametophyte development (Warman et al. 2020a) indicates that the *Zmlarp6c1* transcript (Zm00001d018613_T004) is highest in mature pollen, but largely excluded from the sperm cells isolated from mature pollen. This suggests the transcript is specifically enriched in the pollen vegetative cell, which drives pollen tube germination and growth.

### Supplementary Figure 2

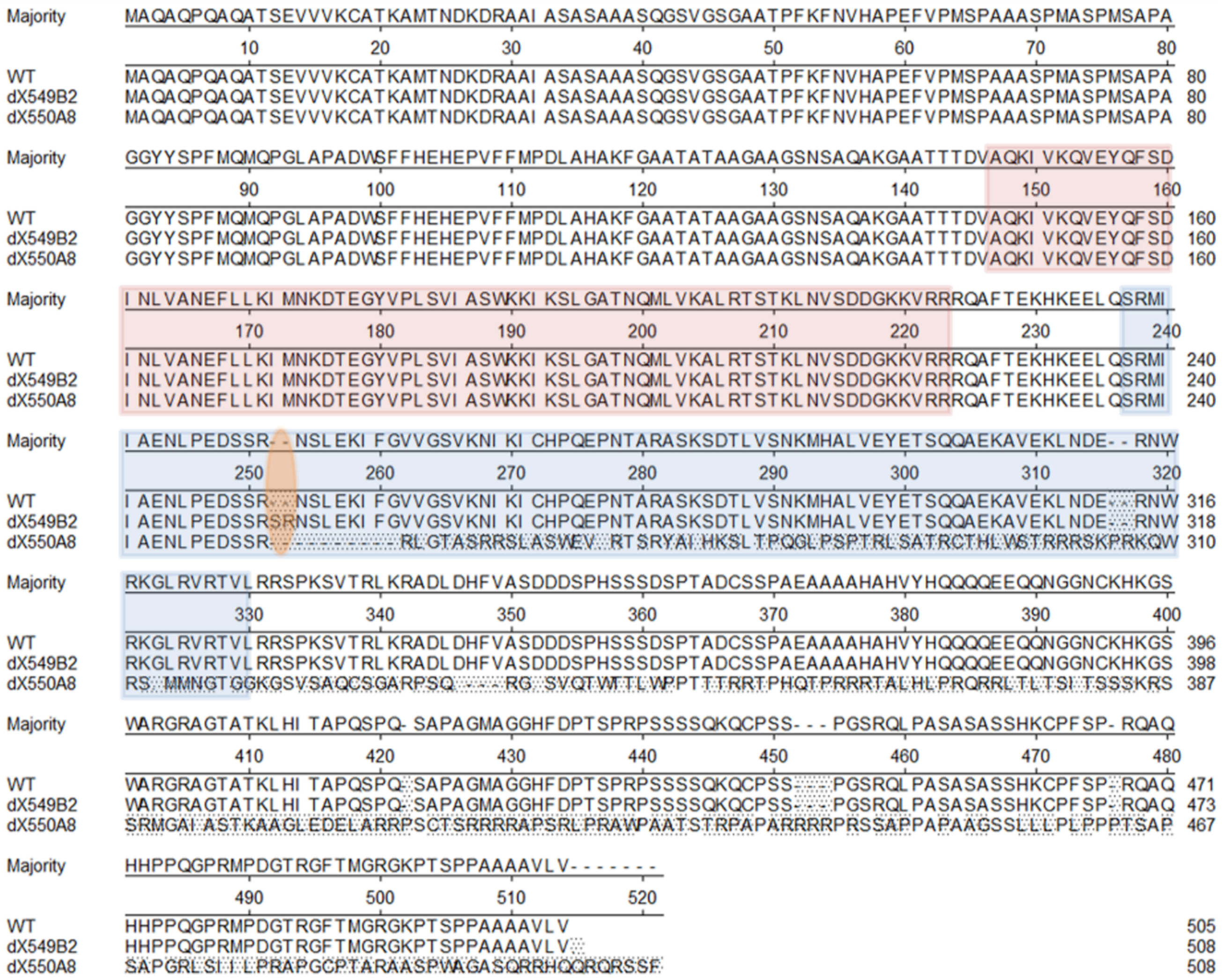


**Supplementary Figure 2 | Alignments of predicted WT, *dX550A8* and *dX549B2* amino acid sequences for ZmLARP6c1.** Red shade, La motif; blue shade, RRM-L; black dot shade, altered residues in derivative alleles; orange oval, two amino acids added by the *dX549B2* six bp footprint at the insertion site.

### Supplementary Figure 3

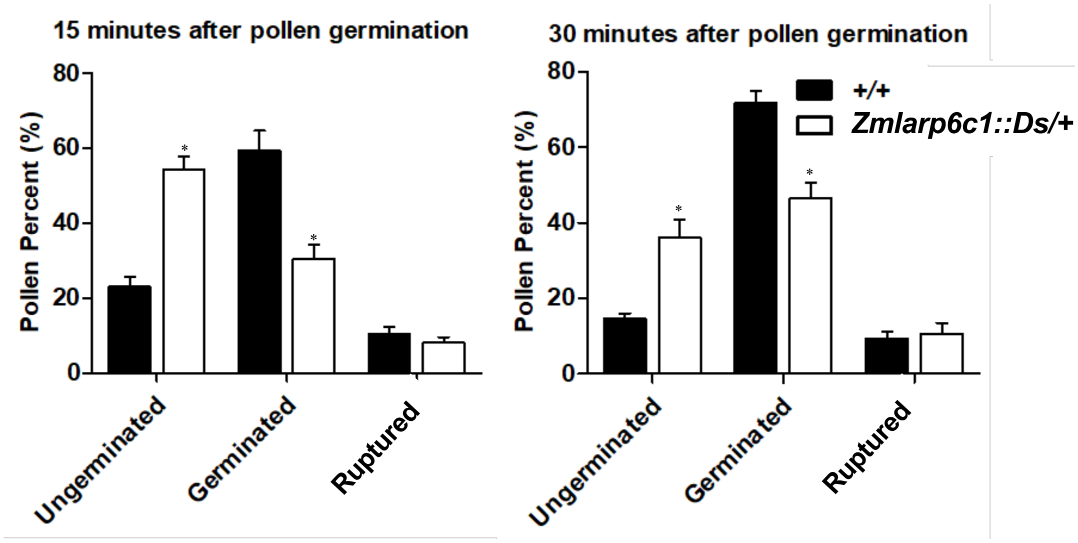


**Supplementary Figure 3 | Pollen germination rate from a heterozygous *Zmlarp6c1::Ds-GFP* plant was lower than from wild-type plant**. Pollen from sibling plants were germinated in pollen growth medium for 15 min or 30 min. Germinated, ungerminated and ruptured pollen percentages were counted (t-test, p-values < 0.01 for categories marked by an asterisk).

### Supplementary Figure 4

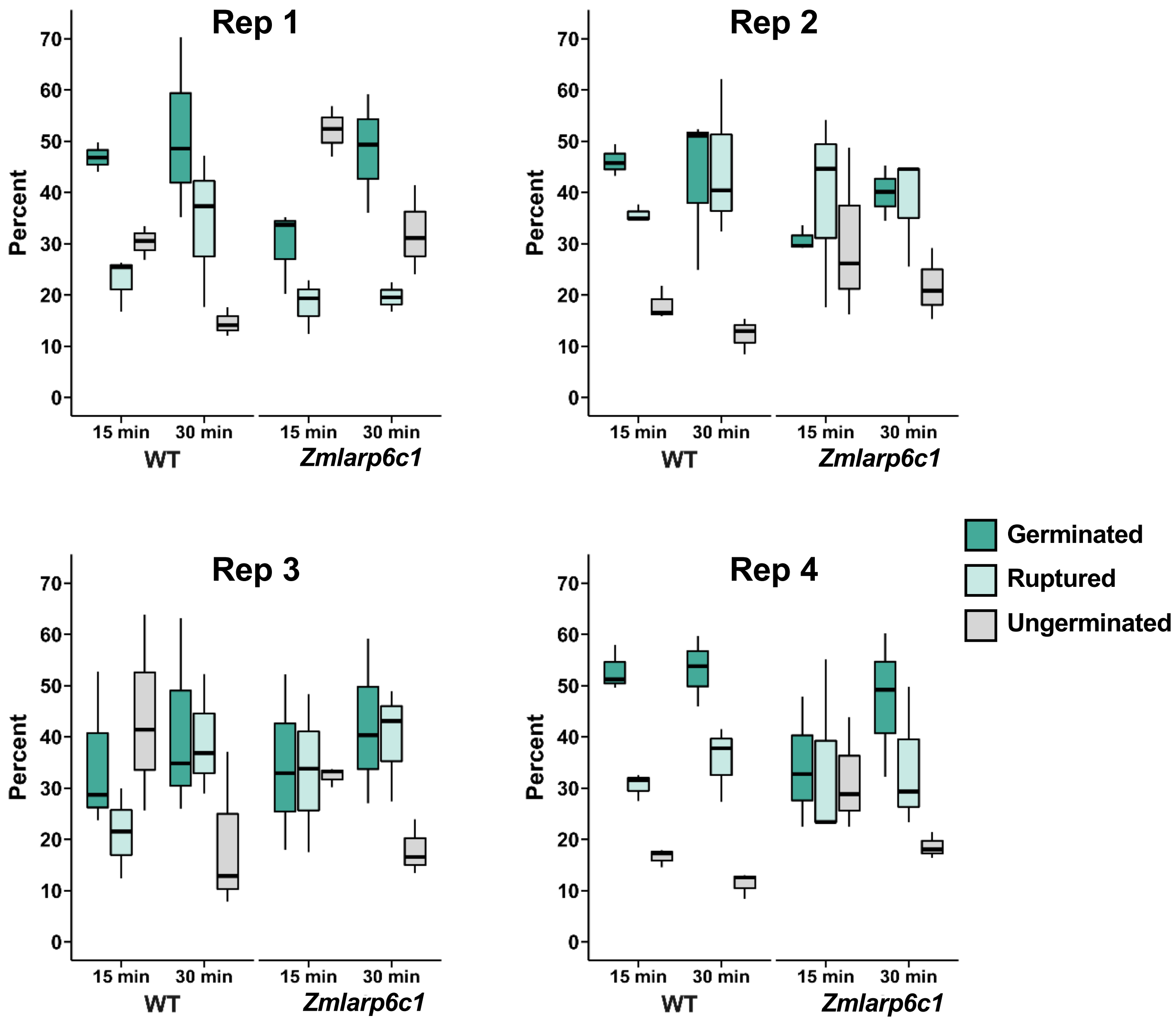


**Supplementary Figure 4 | *Zmlarp6c1::Ds-GFP* and wild-type pollen germination, separated by replicate.** Variance across each replicate is apparent. However, all show a trend toward a smaller percentage of ungerminated pollen grains at 30 min than at 15 min, in both genotypes; and *Zmlarp6c1* pollen associated with a higher fraction of ungerminated pollen than wild type. Modeling the categorical response using a multinomial baseline-category logit model indicates Replicate 2 is a significant factor predictive of a high proportion of Ruptured grains ($p = 0.02059$), whereas Replicate 4 is a significant factor predictive of a low proportion of Ungerminated grains ($p = 0.0134$).

### Supplementary Figure 5

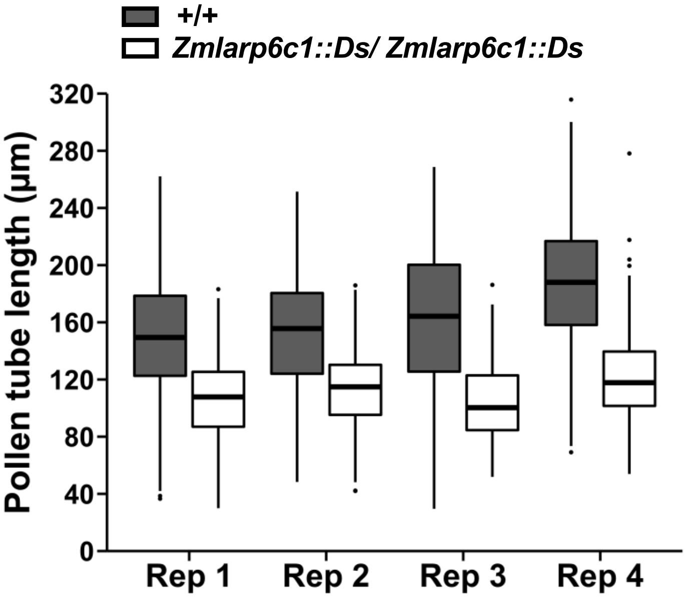


**Supplementary Figure 5 | Pollen from homozygous *Zmlarp6c1::Ds-GFP* plants is associated with shorter pollen tubes.** Pollen from homozygous *Zmlarp6c1::Ds-GFP* and comparator wild-type plants was germinated in pollen growth medium for 30 minutes, imaged, and pollen tube lengths from four replicates were measured. Mutant pollen tubes were significantly shorter than wild type in all replicates (Welch's t-test, all p-values $< 10^{-15}$).
